# NaviNIBS: a comprehensive and open-source software toolbox for neuronavigated noninvasive brain stimulation

**DOI:** 10.1101/2024.11.26.625446

**Authors:** Christopher C. Cline, Lily Forman, Winn Hartford, Jade Truong, Sara Parmigiani, Corey J. Keller

**Author notes:** To whom correspondence should be addressed: Christopher C. Cline. CRediT statement: - Conceptualization: CCC, CJK - Data curation: CCC - Formal analysis: CCC - Funding acquisition: CJK - Investigation: CCC, JT, LF, WH, SP - Project administration: CCC, JT, CJK - Software: CCC - Visualization: CCC - Writing - original draft: CCC, CJK - Writing - review & editing: CCC, LF, WH, JT, SP, CJK.

## Abstract

Image-guided positioning, or neuronavigation, is critical for precise targeting of transcranial magnetic stimulation (TMS) and other noninvasive brain stimulation. However, existing commercial systems have limitations in flexibility and extensibility for research applications. We present new open-source software for neuronavigated non-invasive brain stimulation (NaviNIBS) that provides comprehensive functionality for TMS experiments. NaviNIBS supports imaging data import, target planning, head registration, real-time tool tracking, and integration with robotic positioning and electrophysiology systems. Key features include flexible target specification, support for multiple tracking hardware options, refined head registration techniques, and an extensible addon system. We describe the software architecture, core functionality, characterization of tracking performance, and example applications of NaviNIBS. This software aims to facilitate methodological improvements and novel experimental paradigms in noninvasive brain stimulation research.

## 1 Introduction

Noninvasive neuromodulation techniques have a range of applications, including basic neuroscience research probing brain circuits, clinical diagnostic assessments, and interventional treatments. All common noninvasive neuromodulation methods, such as transcranial magnetic stimulation (TMS), transcranial focused ultrasound (tFUS), and transcranial electrical stimulation (TES), rely on positioning of an external component (or components) on or near an individual’s head, and directing energy (electromagnetic, acoustic, or electrical) into the head, with the goal of modulating activity in the brain. In many cases, the precise positioning of the external component is critical in determining how stimulation energy will be distributed and how the underlying brain tissue will be affected. In this work, we introduce NaviNIBS, a comprehensive open-source system for neuronavigated noninvasive brain stimulation (Cline, 2024a). NaviNIBS provides researchers with a flexible and extensible platform for precisely positioning and tracking neuromodulation devices, integrating advanced registration techniques, real-time visualization, and customizable features to enhance the precision and reproducibility of brain stimulation experiments.

Neuronavigation systems provide a means of monitoring the pose of stimulation components (e.g. a TMS coil) relative to a subject’s head. This pose information is typically presented in relation to a 3D anatomical head model, either constructed from subject-specific imaging data or representative template anatomy. Additional data, such as previously collected functional imaging data (Siebner et al., 2009) or synchronously acquired response data (Sondergaard et al., 2021) may also be incorporated. In this work, we will focus on TMS, but the underlying concepts largely apply to tFUS and TES as well. Use cases for neuronavigation include:

1. informing *initial placement of the TMS coil* based on anatomical features (Yousry et al., 1997; Bungert et al., 2017), common atlas coordinates (Rusjan et al., 2010), previously-collected individual functional and structural connectivity data (Fox et al., 2013; Aydogan et al., 2023), or simulated stimulation effects (Saturnino et al., 2019);
2. measuring the *location of the TMS coil*, selected by other means, for later analysis (Krieg et al., 2015; Rosen et al., 2021);
3. *maintaining consistent coil location* within a session;
4. *returning to the same coil location* across multiple days;
5. *mapping stimulation responses* as a function of TMS coil orientation, such as EMG (van de Ruit et al., 2015; Sondergaard et al., 2021; Weise et al., 2023), EEG (Casula et al., 2022; Gogulski et al., 2024), or behavioral (Tarapore et al., 2013; Krieg et al., 2017) responses.

The most basic features of a neuronavigation system are *subject head tracking*, *subject head registration* to align the virtual model and/or imaging data to real head position, and *coil tracking* to monitor the six degrees of freedom of coil pose relative to the subject’s head. On top of this basic functionality, additional features include image segmentation for head model generation, multimodal data registration, planning of stimulation targets, calibration of tracked tools, variants on head registration procedures, variants on coil pose visualization and quantification, synchronizing of pose samples with stimulation events, integration with electrophysiology response measurements, integration with robotic systems for coil positioning, and more.

Despite the existence of a multitude of commercial and open-source neuronavigation systems and components, there has been no neuronavigation system that is both sufficiently feature-complete to be used in typical TMS studies and extensible for novel neuromodulation research. Commercial neuronavigation systems (Brainsight – Rogue Research, n.d.; Localite: TMS Navigator, n.d.; Nexstim - NBT System, n.d.; visor2 Neuronavigation System, n.d.) are in widespread use in research and clinical settings. Although these typically have the benefit of commercial support and ready-to-use hardware, they are not open source and their limited extensibility can hamper efforts for integration with third-party systems and novel research methods. In the research domain, some open-source neuronavigation systems have been made available (Ambrosini et al., 2018; Souza et al., 2018; Preiswerk et al., 2019) but are not widely used, possibly due to lack of comprehensive software features compared to commercial systems. Additional research efforts have provided demonstrations of and valuable insights into specific neuronavigation approaches and integrations (Leuze et al., 2018; Hassan et al., 2022; Aydogan et al., 2023; Matsuda et al., 2023; Li et al., 2024), but these have largely not been integrated with commercial systems.

Here, we first describe the core architecture and functionality of NaviNIBS, including its advanced head registration techniques, flexible target planning capabilities, and real-time navigation interface. We then detail the software’s extensibility through its addon system, highlighting two key examples: robotic coil positioning and real-time electrophysiological response mapping. Finally, we characterize the tracking performance of NaviNIBS with comparisons to a commercial neuronavigation system. We hope this software will serve as a flexible and powerful platform for researchers, enabling more precise and innovative TMS experiments while fostering collaboration and accelerating development within the brain stimulation community.

## 2 Methods

### 2.1 Software architecture

NaviNIBS is implemented in Python, and uses PyVista (Sullivan and Kaszynski, 2019) and VTK (Schroeder et al., 2006) for 3D visualizations and mesh I/O, Qt via PySide6 (Qt for Python - Qt Wiki, n.d.) and pyqtgraph (PyQtGraph - Scientific Graphics and GUI Library for Python, n.d.) for core GUI components, numpy (Harris et al., 2020) for various computational tasks, pytransform3d (Fabisch, 2019) for manipulation of spatial transforms, and other dependencies mentioned below. Multiprocessing and async operations are used in many parts of the code base to support multiple concurrent or nearly-concurrent operations, focusing on maintaining a responsive GUI during computationally expensive operations.

### 2.2 MRI data

#### 2.2.1 Anatomical data

MRI data, typically T1-weighted anatomical images, can be loaded in NIfTI format; the nibabel library (Brett et al., 2022) is used for reading this data. Multi-axis slice views provide a visualization of the loaded MRI data, as shown in Figure 1A.

**Figure 1:**
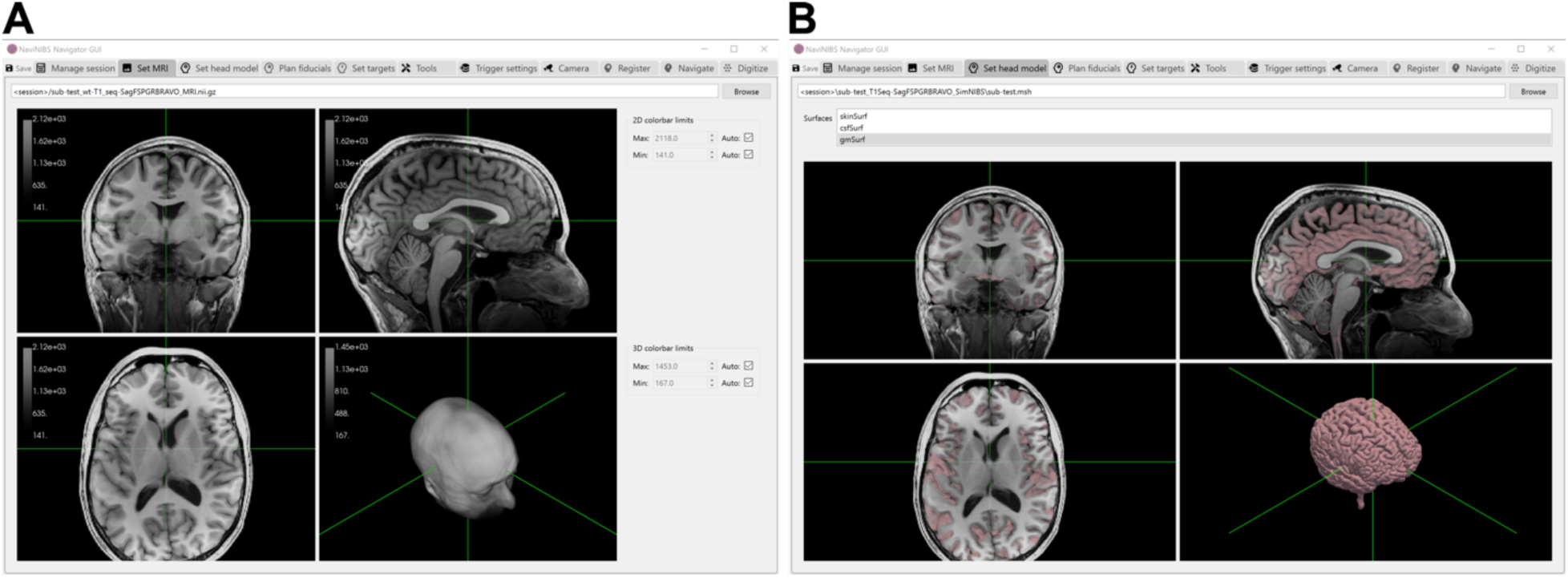
Screenshots of NaviNIBS MRI (A) and head model (B) configuration.

#### 2.2.2 Head model

Unlike many other neuronavigation systems, NaviNIBS does not provide image segmentation functionality to construct a 3D head model from MRI data. Rather than implement our own variant of segmentation procedures, we leverage an existing tool that is increasingly becoming a standard in the field: SimNIBS (Thielscher et al., 2015). Specifically, we use the anatomical head model generated by SimNIBS’ headreco script (Nielsen et al., 2018); support for charm (Puonti et al., 2020) will also be added in a future release. To use a subject-specific head model in NaviNIBS, users can utilize SimNIBS to automatically construct a head model from a T1-weighted anatomical image, optionally supplemented with a T2-weighted image for improved skull segmentation. This head model can then be directly imported into NaviNIBS, as shown in Figure 1B.

#### 2.2.3 Anatomical coordinate systems

The imported T1 MRI data provides a definition of the “native” coordinate system used by various parts of NaviNIBS by which to define relative positions and align other data. In addition, NaviNIBS can also import transforms aligning the subject native space to the group-level Montreal Neuro Institute (MNI) atlas space by linear and nonlinear transformations, generated automatically by SimNIBS or other standard imaging pipelines. These transformations are loaded using nibabel (Brett et al., 2022) and nitransforms (Goncalves et al., 2021).

### 2.3 Tool pose tracking

#### 2.3.1 Tracking hardware

Precise six degrees of freedom (DOF) pose tracking of various objects is critical to a neuronavigation system. This includes tracking of the subject head, TMS coil(s), and a digitizing stylus. A number of possible mechanisms for this tracking exist, including electromagnetic field sensors (Kuehn et al., 2008), 3D laser scanners (Richter et al., 2010), computer-vision based consumer camera systems (Matsuda et al., 2023), and multi-camera optical systems relying on infrared markers. This last option is arguably the most common, with the stereo cameras from Northern Digital Inc. (NDI) being used by at least four commercially available neuronavigation systems. We focus on this optical tracking of infrared markers for NaviNIBS as well, though support for other tracking methods could be incorporated in future releases.

The fundamental principle of tracking with infrared markers is to use at least two cameras, with precisely calibrated relative positions, to estimate positions of markers in 3D space. Using an infrared-sensitive camera and an infrared light source, a marker (in this case, typically an infrared-retroreflective sphere about 10 mm in diameter) can be detected with high contrast from its environment, providing a 2D estimate of marker position with the camera’s field of view. When the marker is visible to at least two cameras, with the camera positions relative to each other known, the 3D position of the marker can be determined. By mounting at least 3 markers with their positions fixed relative to each other as part of a “rigid body”, the 6DOF pose of the entire rigid body, or tool, can be determined. This typically requires that each of at least 3 markers are visible to at least two cameras at any given time.

NaviNIBS supports connecting to NDI Polaris stereo cameras (NDI, Ontario, Canada), with rigid body marker arrangements specified within standard NDI .rom files. NDI Polaris devices are stereo cameras, with a shared housing containing two individual camera units that have their relative positions factory-calibrated; they are typically used as a single device. To maintain pose tracking, this setup requires keeping at least 3 markers on each rigid body visible to the stereo camera.

NaviNIBS also supports a multi-camera localization setup using OptiTrack motion capture cameras (NaturalPoint, Corvallis, OR). While OptiTrack offers monolithic stereo cameras similar in form factor to the NDI Polaris devices, they also offer more flexible and modular systems using reconfigurable arrays of multiple single cameras. These cameras can be mounted around an experiment room, providing more resilience to occlusion of any one camera’s view and potentially greater pose estimate accuracy.

For prototyping purposes, NaviNIBS can also receive tracking information from consumer virtual reality hardware, using infrared-based base stations or inside-out camera views fused with inertial measurement unit data to estimate poses of controllers and headsets. However, with the hardware tested, this was not of sufficient accuracy for typical brain stimulation research, and primarily serves as a low cost entry point for basic software exploration.

#### 2.3.2 Tracking software

Within NaviNIBS, tool pose data is received via the IGTLink (Tokuda et al., 2009) protocol, using the pyigtl library (Lasso, 2023). Using the Plus Toolkit (Lasso et al., 2014), data from either NDI and OptiTrack systems can be streamed via IGTLink to NaviNIBS. The Plus Toolkit also provides similar support for streaming tracking data from a number of other vendors’ devices, but other systems have not yet been tested with NaviNIBS. Additionally, we have implemented functionality for simulating tool positions, allowing GUI- and script-based manipulation of simulated tools for testing purposes. NaviNIBS provides indicators of which tools are visible at any given time, and a 3D visualization of visible tools in the tracked space.

#### 2.3.3 Tool definition

Any number of tools can be configured in NaviNIBS (Figure 2A). Tools assigned to specific types (subject tracker, pointer, coil) will be automatically used for those roles elsewhere in the software. Each tool can be hidden or deactivated when not being used for a portion of an experiment. NaviNIBS supports tracking of multiple coil tools, with the topmost active coil used by default for primary neuronavigation visualizations and reported error metrics. Each tool can have an associated key to associate it with the real-time tracking data stream. Alternatively, tools can be initialized to a fixed location in world space (e.g. a visualization of the tracking camera’s field-of-view) or relative to another tool (e.g. one segment of a robotic arm defined relative to another).

**Figure 2:**
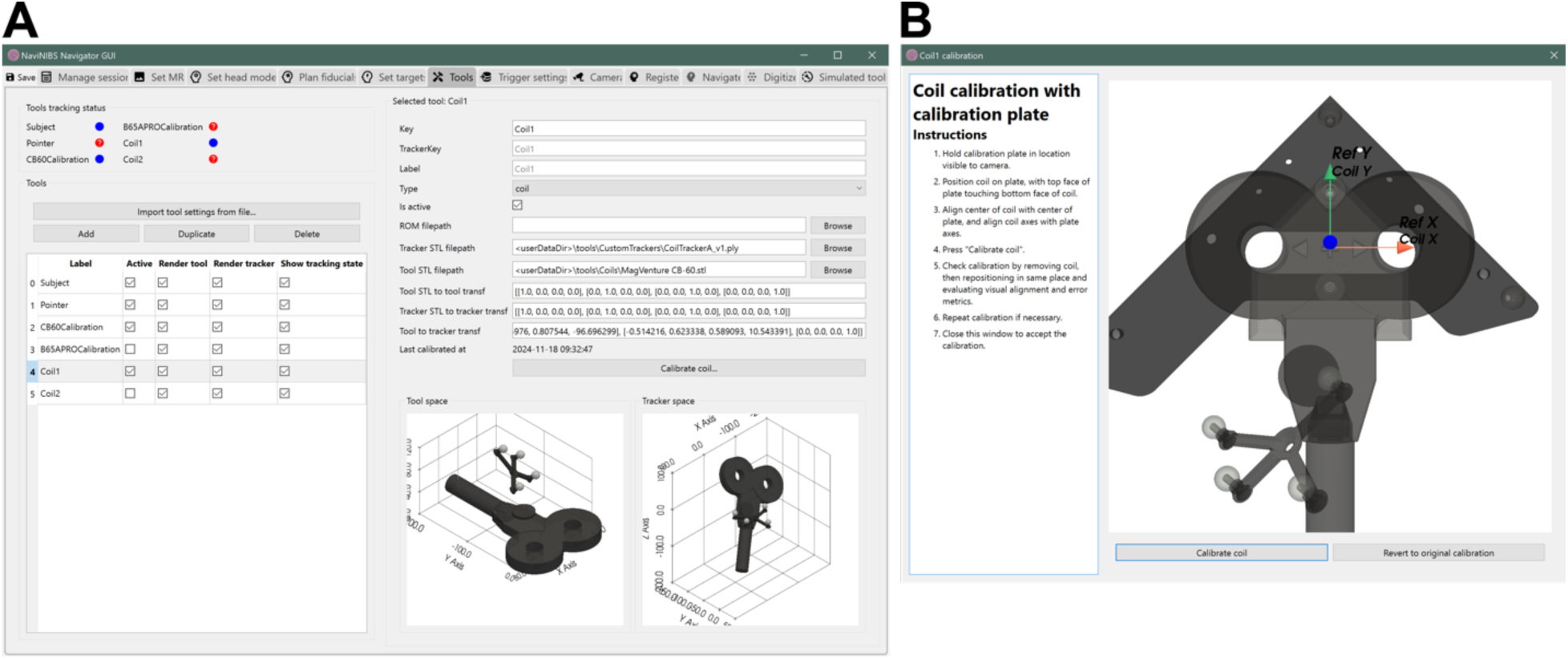
Screenshots of tool configuration (A) and coil calibration (B) in NaviNIBS

#### 2.3.4 Tool calibration

The optical tracking hardware used by NaviNIBS does not directly measure the pose of every tracked tool (e.g. TMS coil), but instead measures the positions of retroreflective infrared markers affixed to the rigid body. For neuronavigation, we must convert from the reported position of these markers to the position of the tool of interest.

For hardware components such as a conventional TMS coil, it is typical to clamp a single rigid body attachment with reflective markers, or “coil tracker”, onto the coil handle. If this coil tracker moves relative to the coil, marker positions relative to the coil must be recalibrated. NaviNIBS supports performing this calibration using a calibration plate – typically a rigid metal plate with predefined marker mounts and a coordinate origin to which the TMS coil is aligned. 3D-printed jigs specialized for the coil being calibrated can help with properly aligning to the calibration plate. The spatial transformation from the coil tracker to the coil can be determined using the spatial transformation from the calibration plate markers to the coil tracker when the coil itself is properly aligned with the plate origin. See Figure 2B for a visualization of this process in NaviNIBS.

A tracked stylus tool is typically used to digitize points, as described later below. Similar to the coil calibration plate, the stylus may be a rigid metal object with predefined marker mounts. If the exact marker locations relative to the stylus tip are known, this information can be predefined and not require any calibration. However, if there is later any physical change (e.g. slight bending of the stylus tip, causing displacement of the tip relative to the markers), a stylus may also need to be recalibrated. NaviNIBS can perform stylus calibration using an endpoint pivot procedure, in which a static location such as a small (<1 mm) marked point on a rigid table is selected, the stylus tip is held on this point, and then pivoted around the point as multiple pose samples are acquired. NaviNIBS then calculates a least squares estimate of the 3D offset between the reported stylus tip and the pivot point, providing this offset as a correction to obtain the true stylus tip location in future pose estimates. This endpoint pivot calibration does not resolve angle mismatches, such as differences reported vs true stylus shaft angle, but precise endpoint location is usually the only relevant information needed for most neuronavigation stylus use cases. If full calibration is required, a calibration plate may be used to calibrate the stylus as well.

Using a stylus and known locations on a physical TMS coil, it is possible to calibrate the orientation of a coil tracker relative to a coil without a calibration plate, based on relative positions of the stylus and coil tracker when pointing to three or more of these known locations. Support for this stylus-based coil calibration will be added to NaviNIBS in future releases.

### 2.4 Head registration

Subject head position is typically determined by tracking a single rigid body attachment with reflective markers, or “head tracker” affixed to the subject’s head. Similar to tool calibration above, the orientation of this head tracker relative to NaviNIBS’ model of the subject’s head must be calibrated. This calibration procedure is referred to as head registration, and can consist of a number of steps focused on maximizing accuracy and precision of this alignment. NaviNIBS includes support for multiple registration approaches, including those also implemented by common commercial neuronavigation systems, in addition to some novel procedures that may improve re-registration speed and precision. An example screenshot of registration in NaviNIBS is shown in Figure 3B, and the sections below describe each of these components in more detail.

**Figure 3:**
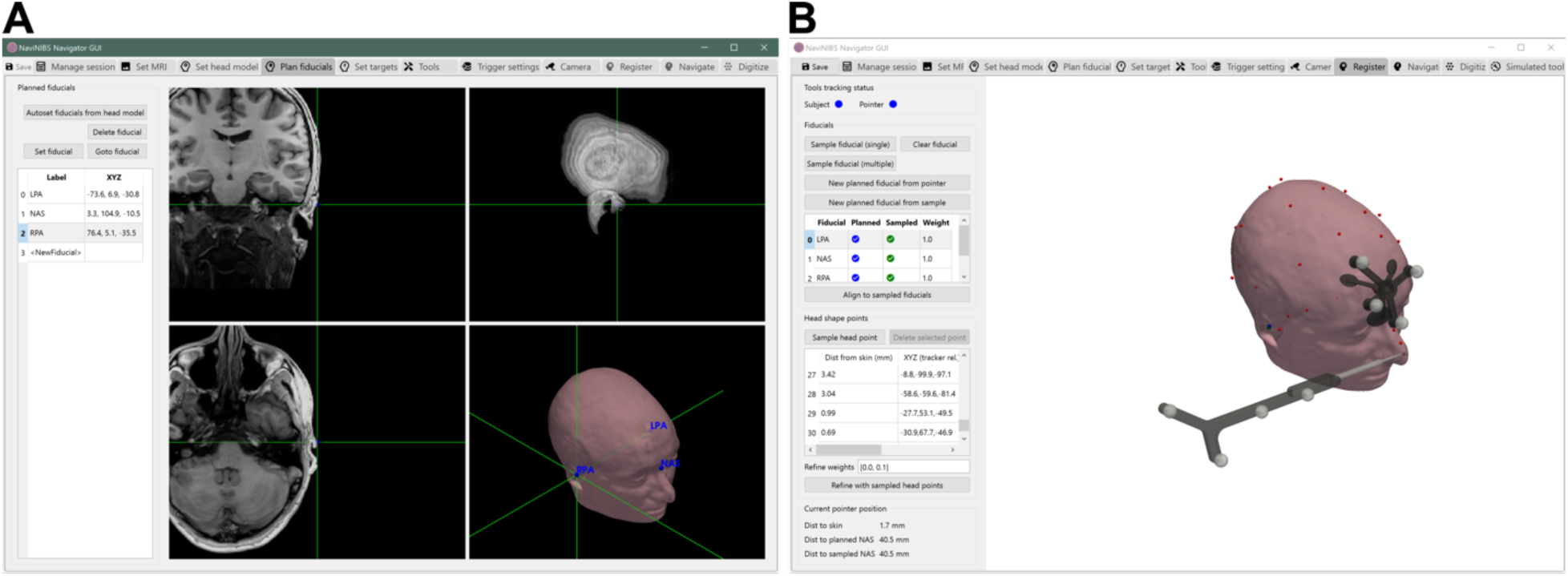
Screenshots of fiducial planning (A) and head registration (B) in NaviNIBS.

#### 2.4.1 Fiducial planning

To align a virtual head model to the subject’s head in real space, a set of common reference points are needed. Typically, a set of anatomical fiducials are used: the most commonly used locations are the left and right preauricular points (LPA and RPA, respectively) and the nasion (NAS). In theory, any point may be used that can be identified both in the subject’s MRI data and pointed to with the physical stylus on the head. However, care should be taken to choose points that are not on soft tissue that may have been displaced during MRI scanning or that may shift during registration. Additionally, the set of fiducials should be spread out as far as possible and not be close to co-linear. For example, using just the left and right inner and outer canthus points without any other points would not produce a reliable registration due to fiducial colinearity.

Planned fiducial locations can be set manually in NaviNIBS by selecting coordinates in orthogonal MRI slice views, or by right clicking directly on the skin surface model. Fiducial locations auto-generated by SimNIBS can also be automatically imported, though these auto-generated locations may need to be adjusted for better fiducial placement (e.g. to a more identifiable location near the pre-auricular points, or away from deflected soft tissue). See Figure 3A for a visualization of fiducial planning.

#### 2.4.2 Initial fiducial registration

Initial head registration is performed by affixing the head tracker to the subject’s head and then pointing with the tracked stylus to each of the planned fiducial locations. A transformation aligning the sampled fiducial locations (relative to the head tracker) and the planned fiducial locations (in the MRI native space) is estimated using a variant of the Kabsch-Umeyama algorithm (Umeyama, 1991).

Once initial registration is complete, the user may assess the quality of alignment by several means. Planned and sampled fiducial locations are visualized separately – large discrepancies between these may indicate an issue with pointing to an incorrect location or having planned a fiducial in a location with soft tissue that was deflected differently during the MRI scan compared to the time of registration. Additionally, NaviNIBS also shows real-time stylus position; by holding the stylus up to various points on the scalp, the user may confirm both visually and quantitatively (based on provided distance metrics) that the distance between true scalp position and head model scalp position is sufficiently small; an example of this is shown in Figure 3B.

#### 2.4.3 Head point refinement

If the planned and sampled fiducial locations are not accurately matched, the initial fiducials-based registration described above may not produce an adequate alignment of true scalp surface to virtual scalp surface. Additional information may be incorporated at this stage to refine the registration, in the form of an unstructured cloud of points sampled on the subject’s scalp. Typically, the user would use the stylus to trace much of the scalp surface and sample something on the order of 100 scalp points broad distributed over the head, avoiding soft tissue like the ears, cheek, and neck. As these points are sampled, NaviNIBS provides distance measures between each point and the scalp, facilitating registration accuracy assessment. Unlike some other neuronavigation systems, NaviNIBS also allows deleting a subset of sampled head points in case of user error without clearing all points.

NaviNIBS can refine the head registration using a weighted iterative closest point algorithm (Glira et al., 2015). Similar to other neuronavigation systems, this shifts the registration to minimize the mismatch between sampled scalp points and the virtual head model scalp surface. Such approaches have been shown to consistently improve head registration accuracy and precision (Nieminen et al., 2022). NaviNIBS provides additional control over this process in the form of separate weighting terms to constrain the translational and rotational components of this refinement. If the user has a high degree of confidence in the fiducial alignment, weights may be set low at this stage to only produce small refinements from the initial registration. If the subject’s head is shaped such that unstructured head point distance minimization doesn’t strongly constrain rotation, the rotation refinement weight specifically may be reduced or zeroed to avoid a rotational refinement of the initial registration, while still allowing for translation to mitigate some systematic shift in planned vs. sampled fiducial locations.

After registration refinement, the head point to scalp distance metrics and fiducial alignment are updated, and the user can use these along with real-time visualization of stylus position to again assess quality of the registration alignment.

#### 2.4.4 Re-registration

To maintain neuronavigation precision, it is critical that the head tracker does not shift on the subject’s head after registration (Nieminen et al., 2022). However, during the long sessions typical during some TMS research studies, it is common for some tracker shift to occur, especially if doing other experimental manipulations on the head (e.g. putting gel in EEG electrodes) or when the subject stands up to take a break. Additionally, in multi-visit studies it is not feasible to place the head tracker in exactly the same position across visits. In such situations, it is necessary to re-register the alignment between the head tracker and subject’s head.

In one approach, re-registration may be performed by simply clearing all previous registration samples (fiducials and head points) and starting from the beginning of the “initial registration” procedure again. This de novo re-registration is what most neuronavigation systems support. This approach should maximize *accuracy* of alignment between the physical head and the virtual head model with each registration, but importantly this does not necessarily maximize *precision* or *reliability* of registration. A random error in sampled vs planned fiducial location, or differences in head point sampling, can contribute to random variations in alignment with each de novo re-registration.

In many TMS studies, precision is more important than accuracy during re-registration. In a study measuring longitudinal effects of stimulation, researchers should typically strive to stimulate in exactly the same location across time periods; if initial registration and refinement produced some inaccuracy between the pre-planned anatomical brain target and the location actually stimulated, that same inaccuracy should be replicated with each re-registration.

NaviNIBS provides several features to specifically maximize precision of re-registration. Described in more detail below, these include: conversion of sampled to planned fiducials, creation of planned fiducials from stylus position, an option to weight some fiducials more than others during alignment, live feedback on stylus-to-fiducial distances, and an option to sample the same fiducial multiple times.

One source of error in registration arises from the user pointing to a location on the head that is different from what was planned in the virtual head model (or MRI data). The user may be very consistent in always pointing to the same fiducial locations with each re-registration, but if those locations differ from the planned fiducials, the mismatch will make it more likely that head point refinement is necessary to obtain a good alignment. Recollecting samples for head point refinement with each re-registration requires additional time and contributes to random variation (i.e. degrades precision) between re-registrations. NaviNIBS addresses this problem by allowing the user to convert previously *sampled* fiducial locations to *planned* fiducials to be used during re-registration. By re-aligning to these previously sampled locations rather than the pre-planned original locations during re-registration, the initial head point refinement from the first registration can be reused without resampling head points, improving registration precision.

Choice of planned fiducial locations are constrained by availability of externally identifiable landmarks on the subject’s head. Typically, this means using points near the eyes, ears, and nose. These points are all biased toward the extreme anterior and lateral regions of the head. With TMS, we typically care most about targeting precision in more central and medial regions of the head. Since the fiducials are far from these stimulated regions, small errors in fiducial location measurement can contribute to larger errors in stimulation target alignment (Nieminen et al., 2022). As an extreme example: if just registering with fiducials at the nasion and left and right outer canthi, a small mismatch in nasion location can produce a very large mismatch in occipital cortex localization.

To address this problem, NaviNIBS supports creation of a new “planned” fiducial sampled from stylus position after initial registration (and refinement). By physically marking a point on the scalp, e.g. with a washable ink pen, one can create a visual reference for a new fiducial location, and sample it with the stylus shortly after initial registration and refinement. Even though this “head reference point” may have no identifiable features in the MRI data, the user can consistently return to that location with the stylus during each re-registration. By choosing a point close to the site of stimulation and using it as a fiducial during re-registration alignment, NaviNIBS can improve alignment precision at the stimulation site.

Related to the idea of specifically minimizing alignment variance near the site of stimulation, it is useful to consider situations when some fiducials may be more “important” than others during registration. NaviNIBS allows specifying a weight for each fiducial, such that some locations are treated with more confidence during registration, using a weighted Kabsch-Umeyama algorithm. In an example with one head reference point weighted several orders of magnitude higher than all other fiducials, this has the effect of pivoting the alignment around that head reference point while best matching the remaining fiducial locations. This weighting can also be useful if some fiducials (e.g. nasion) are more rigid being closer to the skull, compared to others (e.g. preauricular points) being on softer tissue that may deflect differently with each registration, during both initial registration and re-registration.

Another source of error during registration may be caused by the user not consistently pointing to the same location each time the stylus is touched to the head. After initial registration, NaviNIBS provides quantitative feedback with a distance measurement between the stylus location and closest fiducial locations. This allows the user to return to previously sampled fiducials and make sure they are consistently pointing to the same region. NaviNIBS also supports sampling multiple points per fiducial and then averaging their locations for final alignment. Typically the user would position the stylus at the fiducial, record a sample, lift the stylus off and repeat. This gives a simple measure of *individual fiducial repeatability*. Performing multiple samples with multiple stylus orientations around the common tip location also mitigates possible error in stylus geometry or tracking camera distortion, though these are best addressed by other means, such as recalibrating the stylus endpoint location and camera system.

**Table 1:** Summary of re-registration methods.

| Registration Method | Key Features | Advantages | Limitations | Best Use Cases |
| --- | --- | --- | --- | --- |
| Planned fiducials only | - Uses anatomical landmarks<br>- Basic point-to-point alignment | - Quick to perform<br>- Widely used standard<br>- Minimal setup required | - Limited accuracy<br>- Sensitive to landmark identification errors<br>- No refinement | - Initial rough alignment<br>- Short procedures<br>- When speed is priority |
| Full registration with refinement | - Fiducial registration + scalp point sampling<br>- Iterative closest point optimization | - Improved overall accuracy<br>- Compensates for fiducial misregistration errors | - Time-consuming<br>- Requires resampling for each registration<br>- Variable results | - When high accuracy needed<br>- Single-session studies<br>- Initial registration |
| Previously sampled fiducials | - Converts initial samples to new reference points<br>- Maintains consistency across sessions | - High precision<br>- Faster than re-registration with scalp refinement | - Depends on quality of initial registration | - Long duration or multi-session studies<br>- When precision is important |
| Previously sampled fiducials and head reference | - Uses weighted reference point near stimulation site<br>- Combines with sampled fiducials | - Highest precision near target<br>- Faster than re-registration with scalp refinement | - Sensitive to reference point sampling | - Long duration or multi-session studies<br>- When precision is critical |

#### 2.4.5 Recommended registration workflow

With the various registration features described above, the workflow for registration with NaviNIBS may differ in some key ways from existing common neuronavigation procedures. We outline our complete recommended registration workflow below:

1. **Plan at least 3 fiducial locations**, choosing points based on identifiability rather than any strict definition of anatomical feature (e.g. a point fiducial *near* the preauricular point, but not necessarily at it depending on subject’s ear shape), and with large distance between points, and avoiding colinearity.
2. **Mark planned fiducial locations on the subject’s head** with a washable ink pen or wax pencil.
3. **Sample initial fiducial locations**. Hold the stylus in a fixed orientation (e.g. shaft vertical, markers pointing forward) for all samples to prevent any stylus endpoint miscalibration from affecting relative fiducial locations, and carefully match the stylus tip to the marks on the skin within minimal soft tissue deflection.
4. **Perform initial alignment** to planned fiducial locations.
5. **Check for any obvious issues** in fiducial alignments. Try bringing the stylus back to each fiducial location and verify that you are consistently pointing to the same location (based on reported distance to the corresponding sampled fiducial in NaviNIBS). If necessary, modify planned locations for better identifiability and repeat all steps above or just re-sample locations with the stylus.
6. **Sample head points**. Sample broadly and uniformly across the scalp. Avoid regions not well matched in the virtual head model, like soft tissue near the ears and cheeks, the tip of the nose if it was not contained within the original MRI field of view, and areas with segmentation errors like those that can arise near the eyes with some MRI pulse sequences. Make sure to sample with the stylus tip contacting the scalp surface, not elevated off the head (e.g. not on top of an EEG electrode).
7. **Refine alignment using the head points.** Check the results to make sure the refined fiducial locations and scalp points align reasonably with the head model. If needed, modify refinement weights (e.g. by reducing the rotation refinement weight to prefer the rotational orientation set by the initial fiducials), re-align to initial fiducial locations and refine again using the already-measured head points.
8. **Convert the sampled fiducial locations to new planned fiducial locations.** These will be used during re-registration later.
9. **Create a head reference point.** Mark a location on the scalp, approximately near the primary site of stimulation or the top of the head, using a washable ink pen or wax pencil as above. Hold the stylus at this marked location, and create a new planned fiducial from stylus position. It is critical that this be done soon after initial registration (and refinement), before the head tracker has time to shift on the head. Set the alignment weight for this fiducial to a high value (e.g. 1000, compared to the default 1) to emphasize alignment to this fiducial near the site of stimulation more than than the anatomical fiducials during re-registration later.
10. **Continue with typical study procedures.**
11. **Check for registration misalignment occasionally**. If unsure whether the head tracker has shifted at any point, return to the registration pane and place the stylus at the head reference point and (optionally) at the other fiducial locations. If there is a large mismatch in reported distance between stylus position and these locations, then re-registration is needed.
12. **When re-registering**, use the stylus to sample locations for the converted fiducials and head reference points. Align with these new samples and visually check registration. Head point refinement should not be necessary at this stage.
13. If re-registering **across multiple days**, consider documenting precise marked skin locations of the fiducial locations on the first visit (e.g. with close-up photos of the subject’s ears and marked preauricular fiducial locations) to better replicate these marks across days. Depending on the fidelity of replication of these marks across days, head point refinement may or may not be necessary during follow-up visits.

### 2.5 Target planning

#### 2.5.1 Target specification

Stimulation targets in NaviNIBS are defined by a target coordinate (typically on the cortical surface), an entry coordinate (typically on the scalp), an optional offset from entry point to coil location (e.g. accounting for the thickness of EEG electrodes and/or foam between the scalp and coil), and a handle angle (defining the direction of dominant current flow in a typical figure 8 coil). Alternatively, this can be represented as a target coordinate and a spatial transform from the coil coordinate system to the native coordinate space (i.e. 6DOF orientation of the coil in MRI-space). This convention is used by other tools in the field, such as commercial neuronavigation systems and SimNIBS. Coil coordinate system conventions vary between systems; in NaviNIBS, +Z direction points away from the head when the coil is oriented tangentially to the scalp, the -Y direction is assumed to be the direction of dominant current flow or handle direction of a typical figure-8, and the +X direction is orthogonal to these according to a right-handed coordinate system. See the axis indicator in the coil calibration window (Figure 2B) for an illustration of these directions.

#### 2.5.2 GUI-based target creation

New targets can be created in NaviNIBS by simply clicking on the cortical surface of the virtual head model. From this target coordinate, an entry coordinate is automatically generated that minimizes the distance between target coordinate and the scalp, and a handle angle of 0 degrees from midline is initially set. GUI controls allow for editing the target and entry coordinates directly, adjust the approach angles, handle angle, and additional depth offset from the entry. See Figure 4A for an example. Target coordinates can be defined and displayed in subject native or MNI space, or in any other space for which a spatial transform is loaded into NaviNIBS.

**Figure 4:**
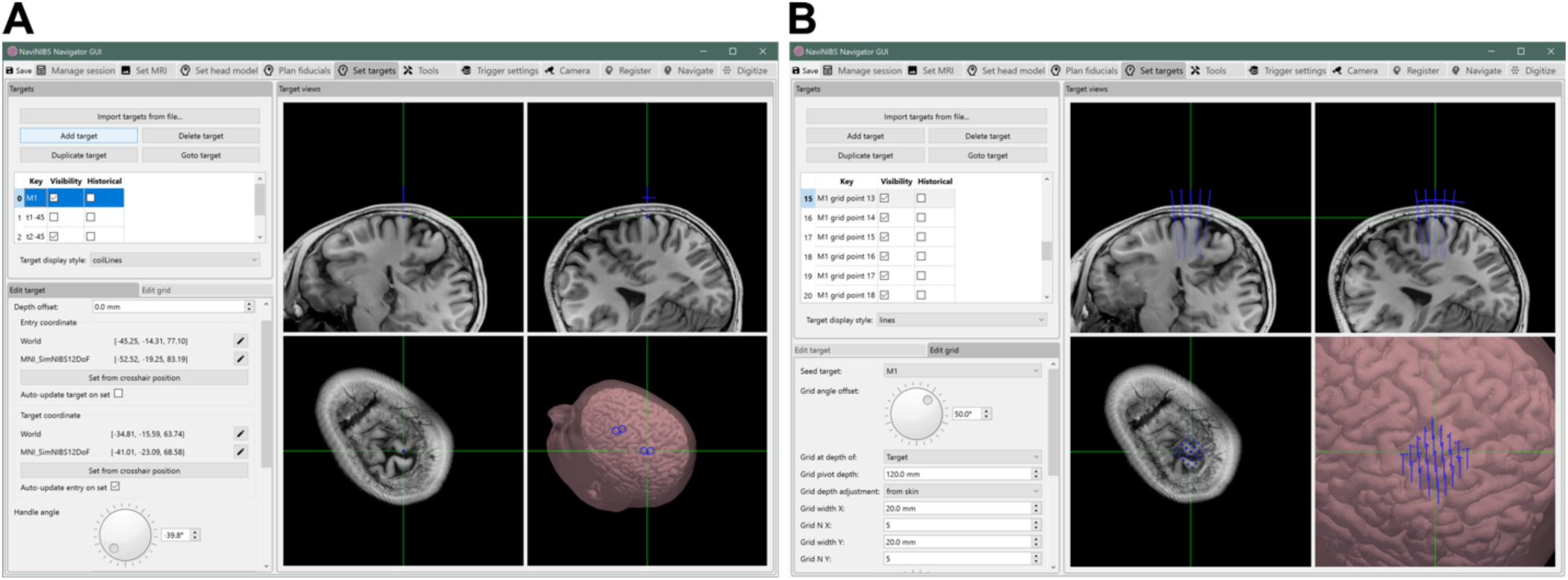
Screenshots of target planning interface in NaviNIBS, for editing a single target (A) and a grid of targets (B)

#### 2.5.4 Target import

Targets can also be imported via files generated by other software tools. Sample MATLAB scripts are available on request to generate suitable .json files, and importers for more direct compatibility with existing tools like SimNIBS will be added in the future.

#### 2.5.3 Target grids

Using an existing target as a seed, grids of multiple targets can be generated. These can be spatial grids, sampling a number of locations at a consistent handle angle in a nearby region, or an angle grid, sampling a number of handle angles pivoted around a shared center target, or a combination of spatial and angle grids. These are defined by a grid span over a direction or angle, number of grid intervals, and optional angle offset in grid alignment from the seed location. See Figure 4B for an example of a generated grid of targets.

### 2.6 Electrode digitization

As part of EEG data analysis, it is common to build a model of EEG signal conduction from the neural sources in the brain to EEG electrodes on the scalp. This benefits from measurements of actual electrode locations on an individual’s head. NaviNIBS provides functionality for digitizing such locations during EEG and TMS-EEG experiments, as depicted in Figure 5. This digitization feature may be useful in other contexts, even if a study does not use EEG: for example, it can be used for measuring the positions of the stimulating electrodes used to produce scalp sensation during sham TMS across subjects or days, or to quantify the precision of head registration, as described in the characterization experiments in section 2.12.

**Figure 5:**
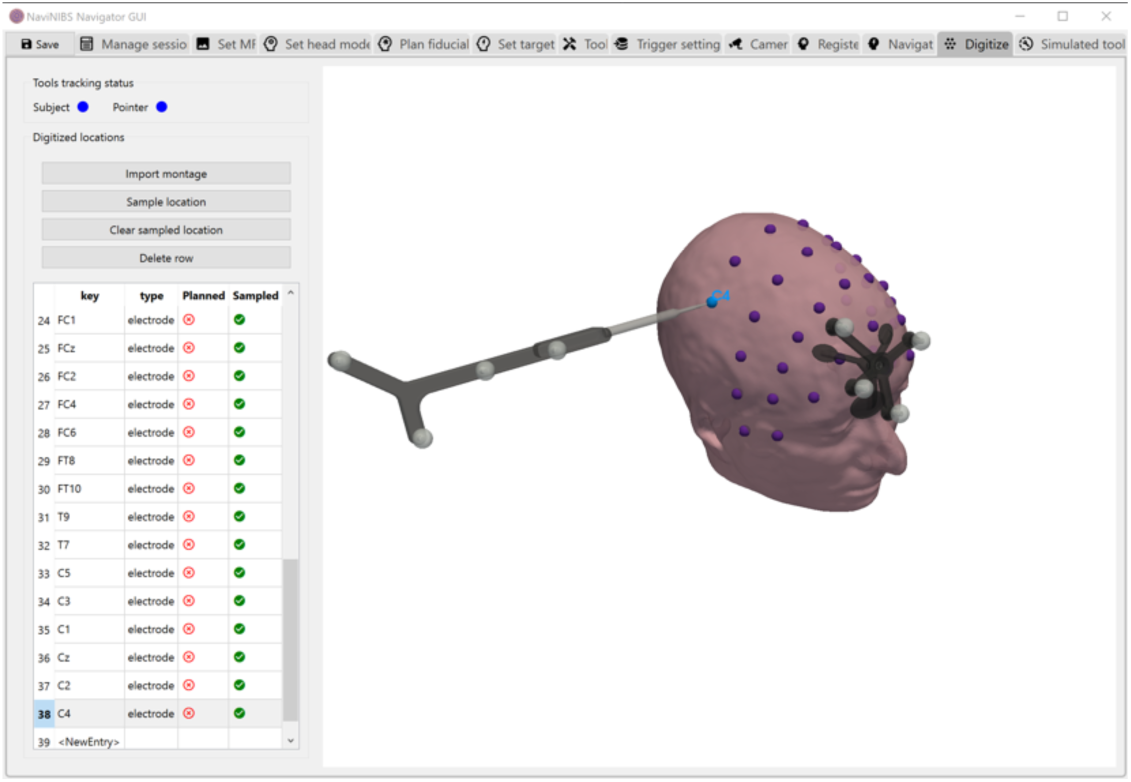
Screenshot of electrode digitization in NaviNIBS

### 2.7 Neuronavigation

With all components of the setup in place, we can consider the core functionality of a neuronavigation system: aligning the stimulating device with a desired target.

#### 2.7.1 Primary navigation visualization

The default visualization style for neuronavigation alignment in NaviNIBS is a set of crosshairs, as depicted in Figure 6b. In this view, in which the camera is looking down at the XY plane of the target coil orientation. The blue crosshairs represent the target, and the green crosshairs represent the current coil pose. The larger diameter crosshairs are at the bottom face of the stimulating coil, and the smaller diameter crosshairs are at the depth of the target in the brain. This stacking of crosshairs allows the user some flexibility in deciding alignment, such as aiming the coil at a common cortical target, even when the depth (Z) axis of the coil is entering the head from a different location. Dominant induced current direction (or handle angle) is indicated by a triangular notch on the crosshairs; other coil angles can be evaluated by keeping the two sets of crosshairs aligned concentrically but rotating the notch around the crosshairs as desired. When the target is misaligned, red lines highlight positioning error between actual and intended target and coil locations, and an orange arc indicates errors in coil angle.

**Figure 6:**
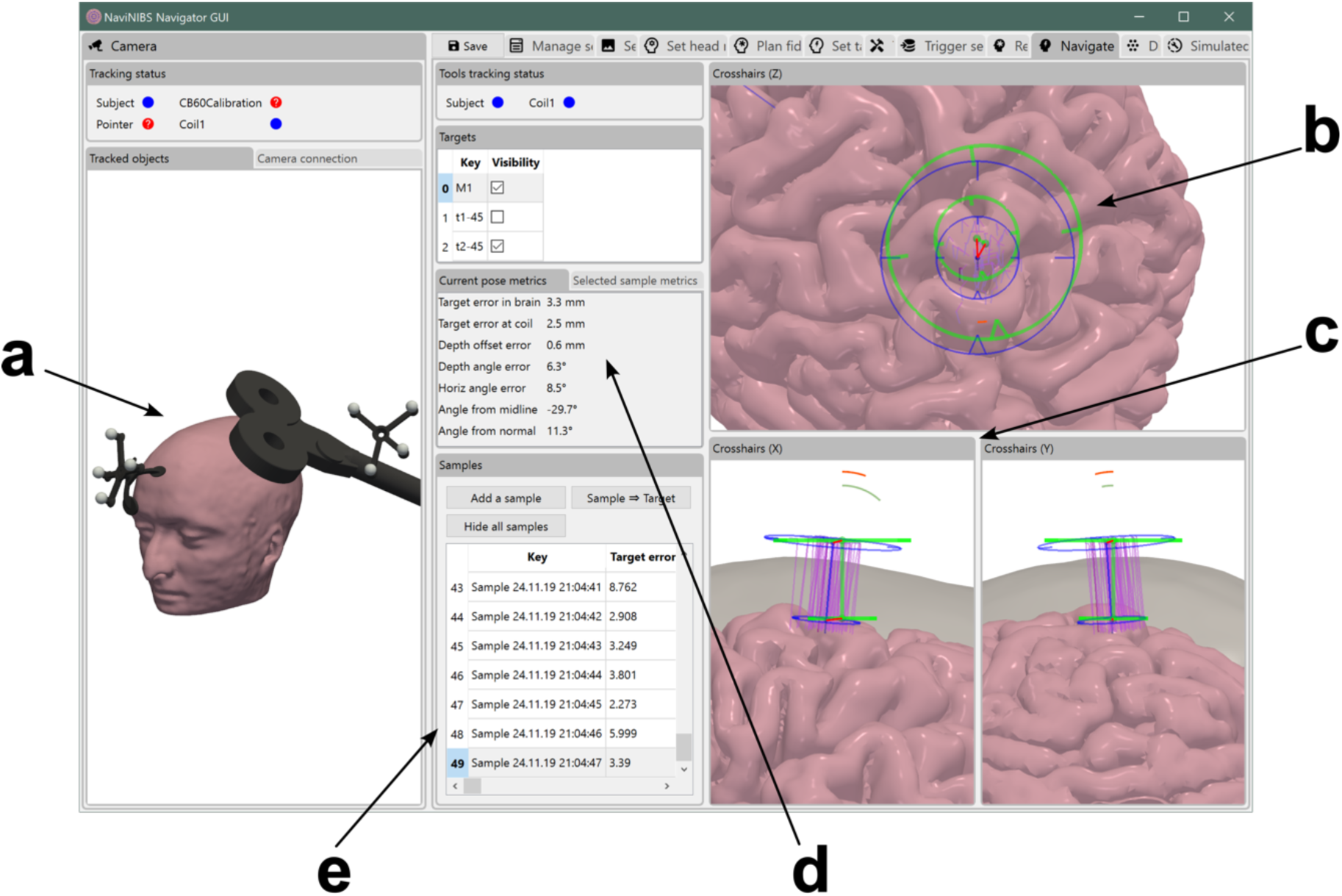
Screenshot of primary NaviNIBS navigation interface, including a “world” view of all tracked devices (a), primary targeting alignment crosshairs (b), orthogonal targeting alignment views (c), positioning metrics (d), and sample history (e).

In perpendicular views depicted in Figure 6c left and right, the camera is looking at the YZ and XZ planes of the current coil orientation, respectively. These views are similar to the Z-axis view, except they allow for more quickly assessing whether the coil is oriented tangential to the scalp at the desired stimulation location. An orange arc indicates entry angle error between the target and actual coil orientation, and a gray arc indicates difference in angle between an auto-estimated “ideal” tangential orientation and the actual coil orientation. The latter indicator is useful to help the TMS technician hold the coil in an ideal orientation while exploring new stimulation sites away from the currently selected target.

In both of these views, the dark purple lines indicate other targets, and the light purple lines indicate recorded samples of past coil positions, as discussed below. These can be hidden to reduce visualization complexity.

#### 2.7.2 Pose metrics

In addition to the visualization described above, NaviNIBS can provide real-time quantitative feedback on coil positioning in the form of pose metrics, as depicted in Figure 6d. These metrics can include absolute measures independent of the current target, such as current angle from midline, but also describe how well aligned the coil is with the desired target. Such metrics can be useful for informing stimulation procedures, such as not starting a stimulation sequence if the target error at the cortical surface is greater than 3 mm.

#### 2.7.3 Coil pose samples

A critical component of many neuronavigation workflows is recording the history of the coil position relative to the subject’s head during stimulation. NaviNIBS provides support for this in the form of coil “samples”, which are defined similarly to targets: a spatial transform from coil-space to MRI-space describes the coil orientation relative to the subject’s head. Additionally, each sample can have an associated target, coil, and other metadata, such as historical pose metrics for each sample. These samples and some metadata fields are depicted in Figure 6e.

New samples can be created manually by clicking the appropriate button in the GUI. More often, however, it is desirable to automatically trigger or create a new sample for each stimulation pulse that is fired in a sporadic single pulse TMS sequence, or every few seconds within a longer repetitive TMS train. NaviNIBS provides support for the triggering from labstreaminglayer (sccn/labstreaminglayer, 2023) streams, a network protocol with support for a wide variety of lab equipment from many manufacturers, including many common EEG and EMG amplifiers, and various open-source software tools. To prevent the sample history from being inundated with too-frequent samples during rapid TMS sequences, a minimum time between consecutive trigger events can be imposed.

A sample can be converted to a new stimulation target to assist with returning the coil back to the same location.

### 2.8 Addon support

As an open-source tool meant to foster new developments in neuronavigation, extensibility is a critical part of NaviNIBS. This is achieved in part through support for “addons”, or modular code components that can be developed separately from the core software and loaded on demand. NaviNIBS provides a number of infrastructure features to support addons integrating with existing functionality.

For example, a NaviNIBS addon may monitor a real-time readout of current coil positioning accuracy and transmit this information out to an LSL stream; source code for this as an example addon is available at github.com/PrecisionNeuroLab/NaviNIBS_LSL_Output. Third-party software could use this LSL stream to decide when to pause a stimulation protocol due to the coil being too far off target.

Two large feature sets implemented with addons are discussed in the sections below.

### 2.9 Robotic coil positioning

In typical TMS experiments, the stimulating coil is either held by hand by a skilled technician, or positioned manually and then clamped into a static coil mount. These approaches can introduce variability in targeting between lab personnel, limit the speed of multi-target mapping procedures, cause difficulties when using heavier stimulating devices like sham-capable shielded liquid-cooled coils, and degrade target tracking fidelity especially in subject populations with greater magnitude and frequency of head movement. One possible solution to these problems is to use a robotic arm to position the coil over the subject’s head (Comeau, 2014; Harquel et al., 2016; Goetz et al., 2019; Dormegny-Jeanjean et al., 2022; Matsuda et al., 2024).

Implemented as an addon, NaviNIBS supports robotic coil positioning with the Axilum TMS-Cobot system (Axilum Robotics, Schiltigheim, France). While some commercial neuronavigation systems also support some basic integration with the Axilum TMS-Cobot, NaviNIBS provides a few key improvements beyond currently available functionality. An example of the visualization provided by NaviNIBS when using the full robotic positioning setup is shown in Figure 7.

**Figure 7:**
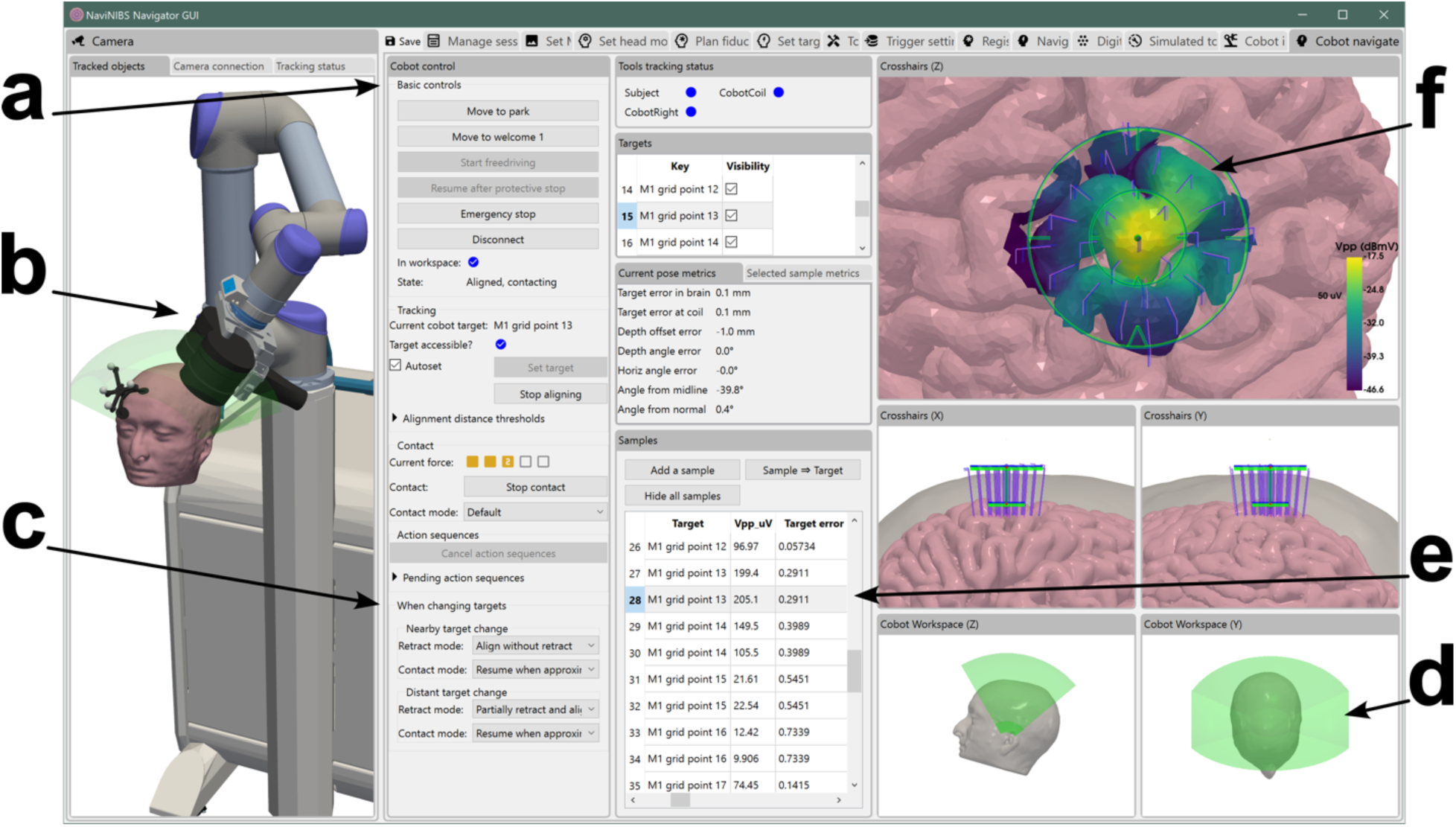
Screenshot of automated motor mapping with NaviNIBS, using Cobot and electrophysiology addons. GUI controls for basic Cobot operations are available (a), as well as more advanced Cobot features like full kinematic visualization (b) and options for reducing time to move between nearby targets (c). The location of the subject within the Cobot’s reachable workspace are visualized (b and d, green volume). With electrophysiology integration, each pulse-triggered coil orientation sample has an associated amplitude response (in this case, simulated MEP amplitude) (d), which can be projected and interpolated onto the brain for mapping visualization (e).

For basic usage, the NaviNIBS-Cobot addon allows sending the coil to a target within the system’s reachable workspace. Head movement is tracked and dynamically compensated, maintaining coil position over the subject’s head, with configurable force sensor sensitivity. Controls are provided for dynamically contacting or retracting from the head, switching targets, and moving to more distant retracted positions (Figure 7a). The position of the subject within the Cobot’s reachable workspace (green volume in Figure 7b and 7d) is visualized in real-time.

With the Cobot system, coil position can be inferred based on optical tracking of the Cobot cart combined with robotic joint angles (Figure 7b), without requiring optical markers on the coil itself. But for greater tracking precision, the NaviNIBS-Cobot addon supports using trackers on the coil and refining Cobot placement based on this information.

The NaviNIBS-Cobot addon dynamically corrects for contact depth offsets, such as can be caused by an EEG electrode in between the scalp and stimulating coil; a similar situation has been observed to cause persistent targeting misalignment in other neuronavigation systems.

When switching between nearby targets, the NaviNIBS-Cobot addon supports several modes of either sliding on the head, partially retracting, or fully retracting the coil before moving to a new target alignment and recontacting the head (Figure 7c). Additionally, optional modes allow for fixing the coil in place (i.e. no longer tracking dynamic head movement) after reaching a target, or for maintaining a non-contact airgap over a given target. This latter functionality may be useful for minimizing the vibration of the coil on the head, at the expense of greater stimulation energy needing to be delivered to the coil for equivalent effect in the brain.

Some of the low-level code for interfacing with the Axilum Cobot is protected under an agreement with Axilum, and is therefore not included in the open-source release of NaviNIBS. However, we have released our higher-level code NaviNIBS-Cobot addon (Cline, 2024b) with an obfuscated low-level module for handling direct communication with the Cobot to be available to the research community.

### 2.10 Electrophysiological response mapping

A common procedure in TMS research is motor mapping, especially to find a location in the hand region of the primary motor cortex (M1) that most efficiently evokes a muscle twitch in the contralateral hand (Sondergaard et al., 2021). This can be done by applying TMS pulses at a number of locations in the vicinity of the hand knob region of the M1 and quantifying muscle activation with bipolar EMG electrodes placed over a particular muscle, such as the first dorsal interosseous (FDI) on the hand.

Using an external software toolbox for real-time signal processing of EMG data, we can extract a motor-evoked potential (MEP) response amplitude, in this case peak to peak potential within 10-45 ms after stimulation, for each pulse. We can then use a NaviNIBS addon to retrieve the quantified response from this external process and encode it in a metadata field associated with each sampled coil location in NaviNIBS (Figure 7e). This response data can then be projected into the cortical surface and interpolated over the nearby cortical mesh, producing a visualization as shown in Figure 7f. Similar visualizations have been shown to reduce motor mapping variability in past work (Kraus and Gharabaghi, 2015). In NaviNIBS, this visualization of the motor hotspot can update in real-time as additional samples are collected at the same or different coil locations. Importantly, this can extend beyond EMG to EEG metrics as well, enabling future applications of closed-loop TMS-EEG with stimulation location guided by observed stimulation responses.

The electrophysiological signal processing toolbox used for this example is under active development in our lab, and describing it in detail is outside of the scope of this paper. However, we plan to release this toolbox and the associated NaviNIBS integration addon in the future. We present it here as a motivating example of the utility and flexibility of NaviNIBS addons.

Combining the robotic coil positioning addon described above with the electrophysiological response mapping addon described here, we can perform fully automated hotspot mapping and similar protocols in NaviNIBS. The results shown in Figure 7f are a simulated result of such a mapping procedure.

### 2.11 Data model

All NaviNIBS session data representing configuration details, head registration, planned targets, sample history, etc. are saved in .json files within a session folder. This format is easily readable in various programming environments, facilitating interoperability with other community pipelines.

For example, sample code is available on request to generate planned targets for loading into NaviNIBS, and to load recorded NaviNIBS session data into MATLAB for retrospective analysis.

### 2.12 Characterizing localization performance

#### 2.12.1 Localization experiments

To validate the registration and tracking performance of NaviNIBS, we conducted a set of experiments quantifying consistency of measured locations on participants’ heads across multiple registration conditions. This study was carried out in accordance with the Declaration of Helsinki and was reviewed and approved by the Stanford University Institutional Review Board. Written and informed consent was obtained from all participants. 6 healthy participants were enrolled.

Two neuronavigation systems were compared: a commercial neuronavigation system (Localite TMS Navigator, Localite, Germany) with an NDI Polaris Vega ST stereo camera (NDI, Ontario, Canada), and NaviNIBS with a separate NDI Polaris Vega ST stereo camera.

At the start of an experiment day, 12 locations were marked on a participant’s head with a washable ink pen: nasion, left and right preauricular points, and 9 locations approximately uniformly spaced over the scalp and aligned with standard 10-10 EEG electrode locations FPz, F3, F4, C5, C6, Cz, Oz, P3, and P4. After each registration condition described below, a tracked stylus was used to record the estimated location of each of these 12 points on the participant’s head using each neuronavigation software’s electrode digitization functionality. An optical head tracker was affixed to the participant’s head, and a “full” registration (alignment to planned fiducials and scalp refinement based on each participant’s T1 MRI) was performed in both the commercial neuronavigation system and NaviNIBS, followed by 12-point measurements (C1 and N1, respectively), with order randomized. These measurements were then repeated to assess short-term reliability (C2 and N2). The head tracker was then removed and re-affixed to the participant’s head in a different orientation, to mimic the need for re-registering in a typical TMS experiment after the head tracker is unintentionally shifted or removed during a break. Several re-registration protocols were then performed (Figure 8A) in randomized order, each followed by re-measurement of the location of each of the 12 marked points on the participant’s head. Full re-registration in both the commercial and NaviNIBS systems (C3 and N3) involved clearing previous registration alignments, aligning to planned fiducial locations, and refining based on newly acquired points on the scalp. Planned-only re-registration (C4 and N4) involved clearing previous registration alignments and aligning to planned fiducial locations without scalp-based refinement. Sampled-only re-registration (N5), only available in NaviNIBS, involved saving the measured fiducial locations from the first registration and re-aligning to these. Samples plus head reference re-registration (N6), also only available in NaviNIBS, involved aligning to the previously measured fiducial locations in addition to a higher-weighted head reference point (located at Cz).

**Figure 8:**
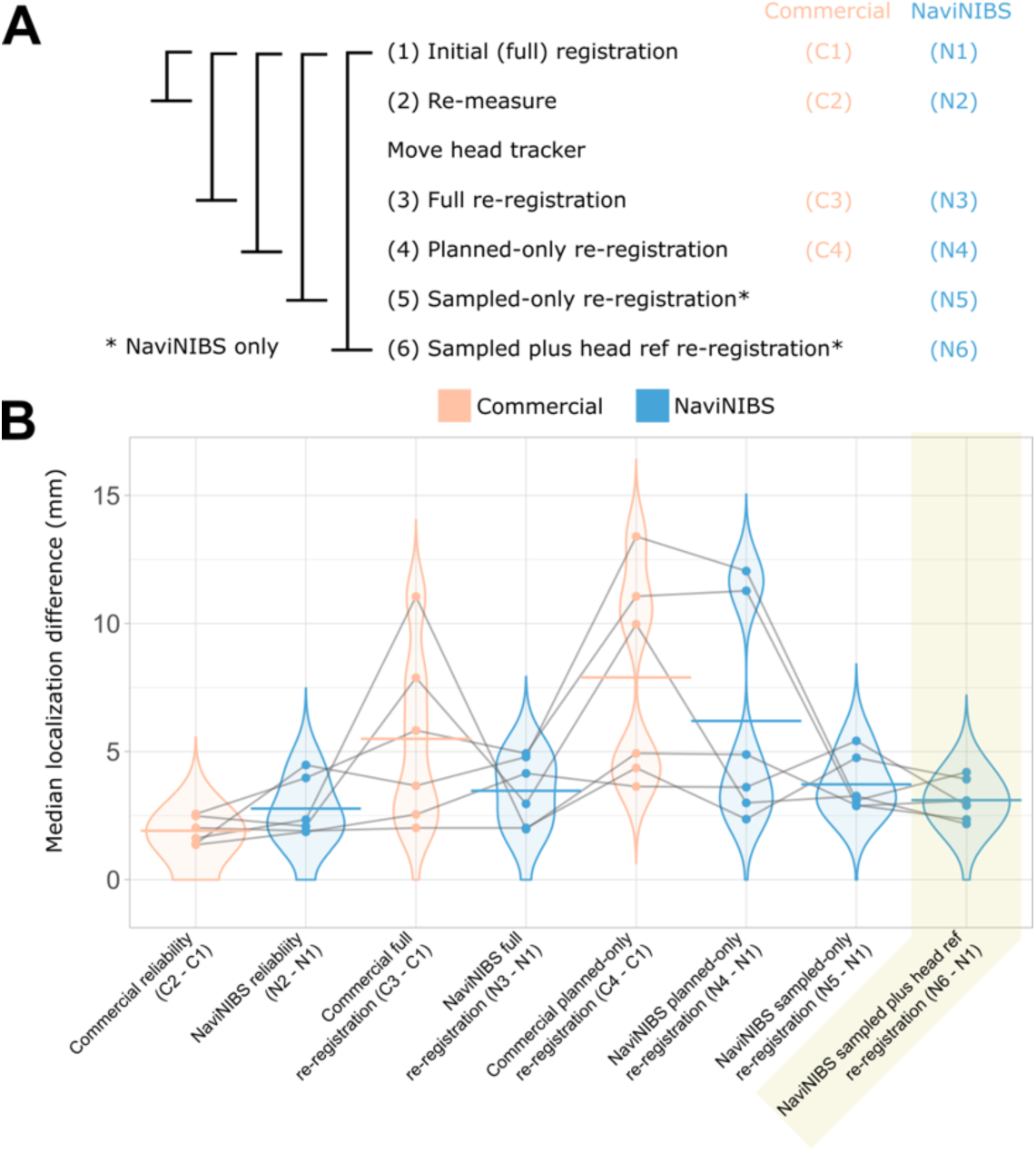
NaviNIBS validation measurement conditions (A) and results (B). In (B), each dot represents median localization difference between two measurement conditions for a single participant; gray lines connect results within a participant; horizontal lines are group means. Condition highlighted in yellow reflects recommended re-registration procedure.

#### 2.12.2 Localization analysis

Localization data was analyzed in MATLAB R2021b (MathWorks, Natick, MA, USA) and R (R Core Team, 2024). Recorded locations were saved in a coordinate space aligned to each participant’s MRI according to the active head registration at the time of measurement. Locations for each successive registration condition were compared to the first set of measurements from the same system, with the difference quantified as the median across the 12 sampled points of the euclidean distance between matching locations, referred to as “localization error” below. Change from measurement (1) to (2) primarily reflects reliability of measurements under identical conditions, with minimal time for head tracker shifts to occur. Change from measurement (1) to (3) reflects reliability of measuring individual points, and of going through the full registration process from scratch. Change from (1) to (4), (5), and (6) similarly represent reliability of measurement and the ability of alternate registration procedures to align with an earlier full registration. Paired t-tests were conducted comparing pairs of localizations for 8 tests of interest: commercial vs NaviNIBS reliability, commercial vs. NaviNIBS full re-registration, commercial planned-only reregistration vs. NaviNIBS planned only, NaviNIBS sampled-only, and NaviNIBS samples plus head reference reregistration, and NaviNIBS sampled-only vs. NaviNIBS samples plus head reference registration, with Bonferroni correction for multiple comparisons.

## 3 Results

### 3.1 Software

Overall, NaviNIBS provides a comprehensive functional neuronavigation solution, and is already in use in other studies in our lab (e.g. (Parmigiani et al., 2024)). The source code for the core NaviNIBS software is at github.com/precisionneurolab/navinibs (Cline, 2024a), and documentation is available at precisionneurolab.github.io/navinibs-docs.

### 3.2 Localization performance

To evaluate NaviNIBS’ tracking accuracy and reliability, we conducted a series of registration experiments across 6 healthy participants. Each participant underwent multiple registration conditions with both NaviNIBS and a commercial system, allowing direct comparison of tracking performance using matching hardware setups. NaviNIBS demonstrated comparable performance to a commercial neuronavigation system across multiple registration conditions when using comparable hardware (Figure 8). Reliability of head point measurements after initial full registrations showed similar localization error for both systems (mean [range] of 1.92 [1.36, 2.57] for C2-C1 vs. 2.78 [1.87, 4.48] mm for N2-N1). Localization error after full re-registration was slightly lower for NaviNIBS (N3-N1, 3.47 [1.97, 4.93] mm) vs. the commercial system (C3-C1, 5.50 [2.02 11.05] mm). The sampled fiducials only (N5) and sampled fiducials plus head reference point methods yielded lower median localization differences (N5-N1, 3.72 [2.88, 5.42] mm; N6-N1, 3.11 [2.18, 4.20] mm) compared to the planned-only re-registration approaches used by both systems (C4-C1, 7.89 [3.64 13.40] mm; N4-N1, 6.20 [2.36, 12.05] mm), presumably due to planned-only approaches not accounting for mismatches between planned and sampled fiducial locations typically captured during scalp refinement. Notably, the sampled plus head reference re-registration method (N6) exhibited the lowest median localization difference of all re-registration conditions tested. Based on paired t-tests, none of these differences were significant after accounting for multiple comparisons.

## 4 Discussion

NaviNIBS represents a significant advance in neuronavigation software for noninvasive brain stimulation research. It provides a comprehensive set of core features for use in typical TMS experiments, and a modular and extensible architecture well-suited for research with novel neuronavigation methods.

The cost of switching to NaviNIBS from another neuronavigation system should be minimal. Integration with SimNIBS (Thielscher et al., 2015; Nielsen et al., 2018) ensures that imaging data is well aligned with other pipelines already typically used in research. Hardware compatibility with NDI Polaris cameras and flexible tool configuration allows the use of tracking hardware from other neuronavigation systems. With parity in core neuronavigation functionality, most workflows from common commercial systems should also be possible with NaviNIBS. Finally, NaviNIBS’ open data structure format makes it straightforward to import pre-planned targets and easily analyze neuronavigation data after a session.

NaviNIBS offers a testbed for improvements in fundamental neuronavigation methods. As an example of this, we have implemented and characterized the performance of several new head registration procedures that can improve targeting precision and reduce registration time in longer sessions or multi-session studies. This includes re-registering to previously sampled fiducials and a head reference point close to the site of stimulation, providing immediate visual and quantitative feedback on registration alignment, and optionally weighting fiducials during alignment and refinement. Such techniques for improving registration precision may be particularly important for longitudinal studies and those investigating subtle effects of stimulation site variations.

Our tracking performance characterization results support the use of NaviNIBS as a viable alternative to commercial neuronavigation systems, while also highlighting potential advantages of its innovative re-registration techniques. Performance in initial registration and re-measurement reliability were comparable. Furthermore, the superior precision demonstrated by NaviNIBS’ sampled fiducials plus head reference point re-registration method suggests a significant improvement over standard re-registration when re-acquiring scalp head points is not feasible. This enhanced precision could be particularly valuable in longitudinal and multi-session TMS studies, where maintaining consistent targeting across multiple registrations is essential, or in subject populations where the subject tracker may be shifted frequently. While these findings are promising, further investigation into the impact of these registration differences on physiological measures and electric fields (e.g. (Nieminen et al., 2022)) would provide additional insights.

An important benefit of NaviNIBS is its open-source implementation and extensibility. By providing a transparent, modifiable platform, it encourages collaboration and innovation within the TMS research community. Researchers can not only use the software but also contribute to its development, potentially accelerating the pace of methodological advancements in brain stimulation research. Code for existing features can be easily reviewed and modified for new experiment needs, and source shared freely to support replication and open scientific review. Entirely new sets of features can be packaged in modular addons, as we have highlighted here with robotic coil positioning and electrophysiological data visualization.

One of our initial motivations for developing NaviNIBS was to have more control of robotic positioning of a TMS coil for automated mapping studies. With the NaviNIBS-Cobot addon, we now have advanced functionality including an API for external algorithms to change the active target, tunable dynamic head movement compensation, rapid switching between nearby targets, more accurate coil tracking, and compensation for EEG electrodes between the coil and head.

We are separately developing a real-time electrophysiological data processing toolbox, and have demonstrated integration with an electrophysiology-focused NaviNIBS addon here. Such functionality can provide a more intuitive interface during common TMS procedures like motor mapping, and enable novel approaches such as real-time EEG-informed target selection in the future. Combined with robotic positioning, we have demonstrated fully automated mapping that has the potential to accelerate and improve the reproducibility of common stimulation targeting procedures.

While we are already using NaviNIBS for neuronavigation in our own TMS-EEG studies, several key limitations should be noted. NaviNIBS in its current form is best suited for use in research pushing the limits beyond existing neuronavigation solutions. In comparison, commercial neuronavigation systems have the advantage of typically being packaged with ready-to-use hardware, mature and stable software, and paid customer support. We are continuing active development of NaviNIBS to improve robustness in core use cases, enhance existing documentation, and implement new features.

One notable feature available in many neuronavigation systems not yet available in NaviNIBS is support for fitting a template head model based on participant scalp measurements and running a neuronavigation session without any participant-specific MRI data. In contrast, NaviNIBS currently requires MRI data to be available. Adding support for neuronavigation with template MRIs is one of the immediate next steps in our future roadmap.

Another feature not yet available but that we plan to implement soon is electric field (e-field) modeling. Specifically, we plan to build on our existing SimNIBS head model integration to also send coil poses to SimNIBS for e-field simulation and then visualize results in the NaviNIBS targeting interface, as well as dynamically calculate metrics like peak e-field magnitude on the cortical surface. This will facilitate more personalized stimulation protocols, supporting experiments that tailor stimulation intensity and location based on estimated field strength.

Overall, NaviNIBS represents a significant step forward in neuronavigation software for noninvasive brain stimulation research. By providing a comprehensive yet extensible platform, we aim to facilitate methodological improvements and enable novel experimental paradigms. As examples of the benefits of this open platform, we have demonstrated new techniques to improve head registration precision, integration with third-party sources of electrophysiology data, and advanced robotic control of TMS coil positioning. With these features, NaviNIBS is positioned as a valuable tool for researchers pushing the boundaries of brain stimulation applications. We look forward to continued development of NaviNIBS as driven by our own experiment requirements, and by requests and contributions from the research community.

